# Topographic somatosensory imagery for real-time fMRI Brain-Computer Interfacing

**DOI:** 10.1101/296640

**Authors:** Amanda Kaas, Rainer Goebel, Giancarlo Valente, Bettina Sorger

## Abstract

Real-time functional magnetic resonance imaging (fMRI) is a promising non-invasive method for brain computer interfaces (BCIs). BCIs translate brain activity into signals that allow communication with the outside world. Visual and motor imagery are often used as information-encoding strategies, but can be challenging if not grounded in recent experience in these modalities, e.g. in patients with locked-in-syndrome (LIS). In contrast, somatosensory imagery might constitute a more suitable information-encoding strategy as somatosensory function is often very robust. Somatosensory imagery has been shown to activate somatotopic cortex, but it has been unclear so far whether it can be reliably detected on a single-trial level and successfully classified according to specific somatosensory imagery content.

Using ultra-high field 7-T fMRI, we show reliable and high-accuracy single-trial decoding of left-foot vs. right-hand somatosensory imagery. Correspondingly, higher decoding accuracies were associated with greater spatial separation of hand and foot decoding-weight patterns in primary somatosensory cortex (S1). Exploiting these novel neuroscientific insights, we developed – and provide a proof of concept for – basic BCI communication by showing that binary (yes/no) answers encoded by somatosensory imagery can be decoded with high accuracy not only offline but also in real-time.

This study demonstrates that body part-specific somatosensory imagery differentially activates somatosensory cortex in a topographically specific manner; evidence which was surprisingly still lacking in the literature. It is also offers a promising novel somatosensory imagery based fMRI-BCI control strategy, with particularly high potential for visually and motor-impaired patients. The strategy could also be transferred to lower MRI field strengths and to mobile functional near-infrared spectroscopy. Finally, given that communication BCIs provide the BCI user with a form of feedback based on their brain signals and can thus be considered as a specific form of neurofeedback, and that repeated use of a BCI has been shown to enhance underlying representations, we expect that the current BCI could also offer an interesting new approach for somatosensory rehabilitation training in the context of stroke and phantom limb pain.

## Introduction

Brain-computer interfacing refers to the use of a signal originating from neural (cerebral) activity to control an external device (Daly & Wolpaw, 2008; van Gerven et al., 2009; Wolpaw, Birbaumer, McFarland, Pfurtscheller, & Vaughan, 2002). While the first BCIs used electrophysiological signals recorded by electrodes penetrating the brain (Fetz, 1969) or by means of electroencephalography (EEG, Farwell & Donchin, 1988), more recently, functional magnetic resonance imaging (fMRI) and functional near-infrared spectroscopy (fNIRS), sensitive to hemodynamic changes induced by active neurons, have also been used in the context of BCI communication and control (Bardin et al., 2011; Chaudhary, Xia, Silvoni, Cohen, & Birbaumer, 2017; Gallegos-Ayala et al., 2014; Lee, Ryu, Jolesz, Cho, & Yoo, 2009; Monti et al., 2010; Naci, Cusack, Jia, & Owen, 2013; Naci & Owen, 2013; Nagels-Coune et al., 2017; Naito et al., 2007; Sorger et al., 2009; Sorger, Kamp, Weiskopf, Peters, & Goebel, 2016; Sorger, Reithler, Dahmen, & Goebel, 2012; Yoo et al., 2004). Compared to fMRI, EEG has a higher temporal resolution, is more mobile and cheaper. However, fMRI has the advantage of providing higher spatial resolution and higher sensitivity to subcortical brain regions. Also, hemodynamic BCIs generally allow for almost instant control, requiring less preparation and training of the subject (Nagels-Coune et al., 2017; Sorger et al., 2012). This means that hemodynamic BCIs can be especially useful for applications in a hospital setting that require high accuracy in a short amount of time and, in the case of fMRI, are used on an incidental basis. Such applications can be found in the realm of state-of-consciousness diagnostics (Naci et al., 2012), communication on important medical issues with ‘locked-in’-patients (Sorger et al., 2012) or auxiliary treatment in psychotherapy or rehabilitation.

With hemodynamic BCIs, it is possible for users to apply an explicit strategy or task to control their brain signal and thereby the BCI output. The requirements for the control strategy vary depending on the intended application. In fMRI-neurofeedback applications, users attempt to reach a certain target brain activation profile by practicing and improving a certain mental strategy based on feedback on their performance. In this case, it is important to establish that the mental strategy is in principle feasible and activates the brain regions that underlie the symptoms that the BCI is intended to improve. For diagnostic or communication applications, on the other hand, there is more flexibility with respect to the applied mental strategies. However, the control strategy should preferably be intuitive and allow almost instant control. It should not tax the possibly limited attentional and cognitive resources of patients (Naci et al., 2012). Ideally, the selected control strategy should be tailored to the patient, relying only on those mental capacities that are preserved (Kaufmann, Holz, & Kubler, 2013), to increase proficiency in BCI control. It is therefore important to develop and test a range of different mental strategies with corresponding experimental and data-analysis protocols to ensure that each patient can be offered the most optimal control strategy.

An especially easy-to-implement category of hemodynamic BCI control strategies is mental imagery, because it does not require any particular stimulation device and can be tailored to the preserved mental capacities of the intended user of the communication BCI. After instructing the participant before the scanning session, start and stop instructions during scanning can be given in any sensory stimulus modality (visual, auditory, tactile), as long as they are technically available and compatible with the user’s faculties. Presently, motor and visual imagery are among the most frequently used mental-imagery strategies in the context of fMRI-BCIs. Both have been shown to generate robust brain activation patterns that can be used as a control signal in the context of fMRI-BCIs for communication (Boly et al., 2007; Naci et al., 2012; Sorger et al., 2012), consciousness diagnostics (Monti et al., 2010) and neurofeedback (Scharnowski, Hutton, Josephs, Weiskopf, & Rees, 2012; Subramanian et al., 2011). Other mental imagery strategies that have been tested in the context of BCI control are spatial navigation (EEG: Cabrera & Dremstrup, 2008; fMRI: Monti et al., 2010), inner speech and mental calculation (fMRI: Sorger et al., 2012; fNIRS: Naito et al., 2007), emotional imagery (fMRI: Johnston et al., 2011; Sulzer et al., 2013), mental drawing (fNIRS: Nagels-Coune et al., 2017), thinking ‘yes’ or ‘no’ (fNIRS: Chaudhary, Xia, Silvoni, Cohen, & Birbaumer, 2017; Gallegos-Ayala et al., 2014) and auditory imagery (EEG: Curran et al., 2004; fMRI: Yoo, Lee, & Choi, 2001).

However, many of the above-mentioned imagery strategies could be suboptimal in patients who suffer impairments in the respective sensory or cognitive modalities. The current understanding is that “the nature of mental representations is formed by our perceptual apparatus and experience” (Schmidt, Ostwald, & Blankenburg, 2014). Therefore, it is important to extend the range of feasible imagery strategies available for operating a BCI in a clinical setting with imagery grounded in modalities in which intended BCI users have vivid memories based on recent experience. In this respect, somatosensory imagery could be an ideal candidate strategy, especially suited for older patients as well as patients suffering from locked-in-syndrome (LIS). The somatosensory modality is relatively well-preserved across the life span compared to, e.g., vision and audition. Also, in ‘locked-in’ patients vision is often impaired due to problems with eye-muscle control and drying of the eye ball related to lack of orbital movement, whereas the somatosensory modality is often one of the few sensory channels that is preserved in the complete ‘locked-in’ state (Murguialday et al., 2011). However, to our knowledge, a somatosensory imagery strategy has not yet been tested for hemodynamic brain-computer interfacing.

Therefore, the goal of the current study was to (1) investigate whether somatosensory imagery is a suitable strategy for operating a hemodynamic BCI, and provide a proof-of-concept that it is suitable for online BCI-based communication; (2) explore to what extent its classification is driven by somatotopic information. For this proof-of-concept study, we exploited the high signal to noise ratio of ultra-high field 7T functional magnetic resonance imaging.

Somatosensory imagery recruits the core imagery-generating network (de Borst & de Gelder, 2017; McNorgan, 2012; Schmidt et al., 2014), but also involves primary and secondary somatosensory cortex (de Borst & de Gelder, 2017; McNorgan, 2012; Newman, Klatzky, Lederman, & Just, 2005; Olivetti Belardinelli et al., 2009; Schmidt et al., 2014; Wise, Frangos, & Komisaruk, 2016; Yoo, Freeman, McCarthy, & Jolesz, 2003). In visual mental imagery, the topographic nature of representations has been shown to be preserved in primary visual cortex, even allowing for decoding of imagery content (Klein et al., 2004; Slotnick, Thompson, & Kosslyn, 2005; Thirion et al., 2006). The primary somatosensory cortex also contains topographic representations (hand: Kolasinski et al., 2016; Nelson & Chen, 2008; Pfannmoller, Greiner, Balasubramanian, & Lotze, 2016; Sanchez-Panchuelo, Francis, Bowtell, & Schluppeck, 2010; Schweizer, Voit, & Frahm, 2008; leg/foot: Akselrod et al., 2017; body: Penfield & Rasmussen, 1950; Zeharia, Hertz, Flash, & Amedi, 2015), but although imagery-induced activation has been shown in S1 and S2, it is unknown whether similar somatotopic activity can be induced using somatosensory mental imagery of the body surface. If somatosensory imagery would induce somatotopically specific activation patterns, this would probably simplify (online) classification, and open up the possibility to use it to activate and train particular subregions of the somatosensory map.

## Methods

### Participants

Ten neurologically healthy participants (six males, mean age = 29 years, sd = 5 years, one left-handed) were recruited among students and staff members of the Faculty of Psychology and Neuroscience at Maastricht University (Maastricht, The Netherlands). Participants were screened for MRI counter-indications and signed an informed consent form prior to their participation. Students received a monetary compensation or credit points after the experiment. The study was conducted in accordance with the Declaration of Helsinki and approved by the ethics committee of the Faculty of Psychology and Neuroscience.

### Procedure and experimental design

Two to five days before the actual scanning took place, subjects participated in a training session in which they performed one or more imagery runs in a mock scanner. The purpose of this training session was to instruct them about the task and possible imagery strategies, and to encourage them to try several imagery strategies and select the one that worked best for them, evoking the most stable and vivid somatosensory experience, as assessed by introspection. Two imagery strategies were suggested as possible starting points. The first strategy was to imagine someone gently touching the inner surface of the hand or foot. The second strategy was to apply techniques from autogenic training, imagining a heavy and warm feeling in the hand or foot. The session lasted at least 30min or until the participant indicated (s)he had found a subjectively effective imagery strategy.

During the scanning session, the participants’ heads were stabilized with foam padding to minimize head movement. They were instructed not to move and to keep their eyes closed during the functional runs. They received auditory instructions indicating when they had to start imagining touch to the right hand (“hand”) and the left foot (“foot”), and when they could stop (“rest”). Each participant completed five (participant 7) to six somatosensory-imagery classifier-training runs. Every classifier-training run contained a pseudorandom sequence of nine left-foot (LF) and nine right-hand (RH) somatosensory mental imagery trials of 18s duration, each followed by a 16, 18 or 20s rest period (figure 1). Seven participants also completed one or more answer-encoding runs in which they answered a binary question using imagined touch to the right hand to encode “yes” and imagined touch to the left foot to encode a “no” answer. Each answer-encoding run included five tactile-imagery trials (encoding the same answer). Timing of the imagery trials and rest intervals was identical to the training runs. If possible, each question was followed by two answer runs. In the first answer run, subjects were asked to encode the actual answer to the question using the appropriate imagery strategy. In the second answer run, they were asked to encode the opposite answer (in order to obtain approximately the same number of “yes” and “no” answers-encoding data). The correct answers to the questions were unknown to the researcher responsible for data analysis until after completion of the offline analyses. In one subject, the answer runs were also analysed online and the results were reported back to the participant right after the run.

**Figure 1:**
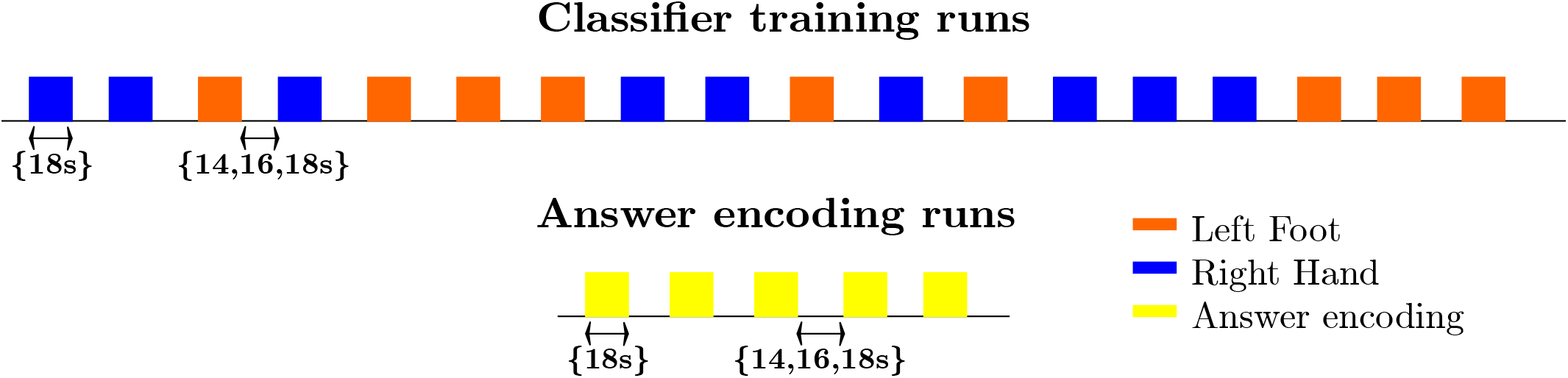
Diagram illustrating the timing and an exemplary pseudorandom temporal order of imagery trials and rest intervals for the classifier-training and answer runs.

After scanning, participants were asked to fill out a questionnaire in which they had to rate the experienced quality of their somatosensory imagery (i.e., whether it was a robust mental experience) and the extent to which it evoked a tactile sensation (i.e., to what extent did they experience it as inducing tactile input coming from the target limb) on a ten-point scale (one being the worst/least applicable, ten being the best/most applicable). They also rated the percentage of time that they were successful at maintaining the imagery and described the particular imagery strategy they used.

All subjects reported successful somatosensory imagery. The type of strategies used ranged from imagining a warm feeling (four subjects), imagining vibratory stimulation to the finger or toe (two subjects), to imagining touch (massage, brush strokes, somatosensory sensation; four subjects; supplementary table 1). On a scale of 1-10, the average score subjects gave for the quality of their hand imagery was 6.9 (sd = 0.65) and of their foot imagery 6.6 (sd = 0.49). The average score in response to the question to what extent they experienced a tactile sensation during imagery was 5.3 (sd = 2.3) for the hand and 4.9 (sd = 1.9) for the foot. The reported average percentage of time that somatosensory imagery worked during the hand imagery epochs was 75% (sd = 4.5%, N=6) and 73% (sd = 8.2%, N=6) for the foot imagery epochs. Paired t-tests did not reveal any significant differences between foot- and hand-related ratings at p<0.05.

### MRI data acquisition

Data were acquired at Maastricht Brain Imaging Centre (Maastricht University) on a 7T scanner (Siemens Healthcare, Germany) with a 32 receiver channel (Nova) head coil.

The scanning session included the acquisition of an anatomical scan (8min) and five to six classifier training runs (12min/run). In seven subjects, there was sufficient time to end the session with one to seven additional answer-encoding runs (1 answer run: 2 subjects, 2 answer runs: 4 subjects, 7 answer runs: 1 subject) in which the participant answered one or more binary questions using somatosensory imagery (4min/run).

For the anatomical scan a Magnetization Prepared 2 Rapid Acquisition Gradient Echoes sequence (MP2RAGE, Marques et al., 2010) was used with 240 slices of 0.7 thickness, in-plane resolution of 0.7mm, 240mm × 240mm field-of-view, TR/TE= 5000/2.47ms, inversion time 1900ms, inversion time 2 2750ms, flip angle 15 degrees, flip angle 2 3 degrees and GRAPPA acceleration factor 3.

In the functional runs, a gradient echo sequence was used with 78 slices of 1.5mm thickness, 1.5mm in-plane resolution, 198 mm × 198 mm field-of view, TR/TE = 2000/26.1ms, 60 degree flip angle, multiband acceleration factor 2 and GRAPPA acceleration factor 3 (340 volumes for the training runs, 96 volumes for the answer runs). In each session, five volumes were acquired in opposite phase encoding direction to be used for (offline) correction of EPI distortions.

## Data analysis

### Preprocessing for offline multivariate analyses

Offline data analysis was performed using Brainvoyager QX 2.8.4 (BrainInnovation, Maastricht, the Netherlands). The anatomical scan was down-sampled to 1mm^3^. The first two volumes of each functional run were discarded. The remaining volumes were adjusted based on slice-scan time, realigned to the first functional image of the session using rigid body transformations, temporally filtered to remove linear trends and nonlinear periodic signals with a frequency below 0.01Hz, corrected for EPI distortions, coregistered to the anatomical scan interpolating to 2mm^3^ isotropic, and normalized to Talairach space. This procedure includes the definition of subject specific anatomical landmarks (AC [anterior commissure], PC [posterior commissure] and the borders of the cerebrum) that are used to rotate each brain in the AC–PC plane followed by piecewise, linear transformations to fit each brain in the common Talairach ‘proportional grid’ system (Talairach & Tournoux, 1988).

The multivariate analyses were limited to the voxels included in anatomical masks. Masks were obtained by creating anatomical regions of interest (ROIs) in Talairach space based on the Jülich probability maps for the primary (PSC3a, PSC3b, PSC2 and PSC2; Geyer, Schleicher, & Zilles, 1999; Geyer, Schormann, Mohlberg, & Zilles, 2000; Grefkes, Geyer, Schormann, Roland, & Zilles, 2001) and secondary somatosensory cortex (SIIOP1, SIIOP2,SIIOP3 and SIIOP4; Eickhoff, Amunts, Mohlberg, & Zilles, 2006; Eickhoff, Schleicher, Zilles, & Amunts, 2006) at different probability levels (see below).

Features (t-values per voxel) were extracted by linearly fitting each voxel within the mask with a design matrix consisting of a constant term (intercept) and a modelled hemodynamic response, within a temporal window of 30s, spanning from one volume (2s) before the auditory cue to 14 volumes (28s) after the auditory cue. The modelled hemodynamic response was obtained by convolving a canonical hemodynamic response function (hrf) with a box-car predictor whose value was one in the volumes corresponding to the imagery blocks, and zero otherwise. The fit was done separately for each trial, and the t-estimates of the hemodynamic model were used as features per voxel. This procedure resulted in 45 (participant 7)/54 feature examples per class (nine per run).

Data were imported in Matlab (R2013b, Mathworks Inc.) using the NeuroElf (v1.0, http://neuroelf.net, Jochen Weber) for feature extraction and Spider toolboxes for multivariate classification. A linear Support Vector Machine (SVM) (Vapnik, 1995) with a constraint parameter C= 1 was used in several leave-run-out cross-validation procedures which differed depending on the research question, and are described in more detail below. Before learning a model, training features were z-scored to ensure the magnitude of weights would be interpretable. To test whether classification accuracies were significantly above chance, 2000 permutations (Golland, Liang, Mukherjee, & Panchenko, 2005) were performed in which condition labels were randomly reassigned to each block in each separate run. The p-value was calculated as the ratio between the number of permutations where the error was lower or equal to the observed one and the total number of permutations (adding one to both numerator and denominator to avoid zero values). Subsequently, proportions correct classifications were computed with edge correction ([number of correctly classified trials – 0.5]/number of trials) and logit transformed before statistically testing the hypotheses of interest using SPSS (IBM SPSS Statistics 24 for Windows, Armonk, NY: IBM Corp.).

### Suitability of somatosensory imagery for BCI

#### Effect of the amount of training data and imagery class

To test the hypothesis that the amount of training data would increase decoding accuracy independent of the imagery class the number of runs used for SVM training was increased from one to five, each time testing on one of the remaining runs, using all possible combinations. Features were extracted from the largest, most inclusive anatomical masks of the primary and secondary somatosensory cortex. To test single-subject significance, permutation tests were performed. The edge corrected and logit transformed classification accuracies were subsequently entered in a two-way repeated-measures ANOVA in with imagery class (2) and number of training runs (5) as within-subject factors. To test whether accuracy increased with the number of training runs, within-subject planned polynomial contrasts were used.

#### Assessment in the context of BCI communication

To test the suitability of the current approach for (online) communication, a simulated realtime classification was performed using Turbo-BrainVoyager (version 3.2, BrainInnovation B.V., Maastricht, the Netherlands) to emulate the procedure that would be performed during the scanning session. In the last participant, a proof-of-concept online communication experiment was performed in which the answer runs were also decoded online during the scanning session using an SVM based on data from all available training runs.

The online and simulated real-time analyses were based on slice data in native resolution (1.5mm^3^), which were 3D-motion corrected using the standard TBV incremental procedure spatially realigning each volume to the first recorded volume by rigid body transformation. The lowest probability (largest) S1 and S2 mask was back-transformed from Talairach space into native space. After spatially smoothing the functional data with a 4 mm full -width -at -half –maximum (FWHM) kernel, t-values were computed by fitting a general linear model with the same predictors as used in the offline procedure, within the same temporal window of 30s, spanning from one volume (2s) before the auditory cue to 14 volumes (28s) after the auditory cue. T-values from the S1 and S2 mask were used as features for SVM training and testing using LIBSVM (2000-2009 Chih-Chung Chang and Chih-Jen Lin).

In the simulated real-time analyses, the effect of the number of training runs was tested by using a classifier based on an increasing number of runs, starting from the run closest to the answer run to emulate the temporal adjacency that is characteristic for the online situation, each time adding a preceding run for each additional level for SVM training, and subsequently testing on the available answer run(s). The resulting edge-corrected and logit transformed accuracies were entered into a repeated-measures ANOVA with number of training runs (1-6) as within factor was used to statistically assess the effect of the amount of training data.

### Assessment of somatotopy

#### Comparing accuracy in primary and secondary somatosensory cortex masks

The effect on decoding accuracy of the anatomical region chosen for feature extraction was evaluated by repeating the cross-validation procedure with five runs (four for subject 7) for training and one run for testing with features extracted from the anatomical masks of the primary (PSC3a, PSC3b, PSC2 and PSC2) and secondary (SIIOP1, SIIOP2, SIIOP3 and SIIOP4) somatosensory cortex at different probability levels (Eickhoff, Amunts, et al., 2006; Eickhoff, Schleicher, et al., 2006; Geyer et al., 1999; Geyer et al., 2000; Grefkes et al., 2001). The probability level (a percentage) of a mask for a particular anatomical area indicates that all voxels in that mask were classified as part of that anatomical area in at least the corresponding percentage of subjects in the database used to construct the anatomical probability map. For example, an anatomical mask with a probability level of 30% contains voxels that were classified as belonging to that particular anatomical region in at least 30% of the subjects in the database used to construct these anatomical probability maps. The higher the probability level, the smaller the mask size (supplementary table 2). To test single-subject significance, permutation tests were performed.

Accuracy values were entered in a two-way repeated-measures ANOVA, with anatomical region (2, S1 or S2) and probability level (8, 10-80%) as within-subject factors, to test whether there was a difference between S1 and S2 in terms of decoding accuracy, and whether this was possibly affected by the probability level, which was further assessed using within-subject planned polynomial contrasts. For the ANOVA, only probability levels of 80% or lower were taken into account which corresponded to masks with more than 500 voxels, to ensure stable classification results.

#### Assessment of somatotopy in discriminative SVM weights

The extent to which the classification was based on somatotopically different weight patterns in primary somatosensory cortex was assessed using the average discriminative weights across the different training splits. Discriminative weights were extracted from single-subject maps (obtained when testing the overall single-subject classification accuracies) that represented the average weights across the different training splits, using SVM classifiers trained on features from the most inclusive anatomical mask, with all possible combinations of 4/5 runs for training. Because the representations of hand and foot are most clearly separated on x-axis (left-to right), we computed the subject-specific median x-coordinates for the 200 largest positive (foot imagery) and 200 largest negative weights (hand imagery), and compared them using a Wilcoxon signed rank test. Finally, the Spearman rank correlation was computed between the subject-specific differences in median x-coordinates for foot and hand and the leave-one-run-out classification accuracy

## Results

Ten healthy participants were instructed by auditory cues to perform alternating somatosensory imagery of the hand or foot (figure 1) while ultra-high field 7T fMRI data were collected. Each participant completed five (participant 7) to six somatosensory-imagery classifier-training runs. Seven participants also completed one or more answer-encoding runs in which they answered a binary question using imagined somatosensory stimulation to the right hand to encode “yes” and imagined somatosensory stimulation to the left foot to encode a “no” answer.

### Suitability of somatosensory imagery for BCI

After establishing that all participants subjectively experienced somatosensory imagery (see methods section), we went on to test the suitability of this strategy to induce activation that can be reliably decoded for the purpose for BCI communication. We hypothesized that, using a machine-learning algorithm, the contents of somatosensory hand or foot imagery can be accurately decoded on the single-trial level with similar accuracy for each imagery class, with accuracy increasing when increasing numbers of runs are used to train the classifier (hypothesis 1a). Our second hypothesis (1b) was that somatosensory imagery would yield accurate decoding of answer runs in simulated real-time as well as during a proof-of-concept online brain-based communication session.

Testing the first hypothesis (1a), we found that significant above-chance classification accuracy (p<0.05, based on permutation testing) was achieved in eight out of ten subjects with one training run, and for all subjects for two or more trainings runs (figure 2). The average proportion correct classifications increased with the number of runs included for training (figure 3), up to 0.82 (sd= 0.12) when using five runs (four in one subject) for training and one run for testing. The effect of imagery class and the amount of training runs was tested in an imagery class (2, foot or hand) by training runs (5) repeated measures ANOVA on the edge corrected and logit transformed accuracy data. This revealed a significant effect of the number of training runs on the decoding accuracy (F(1.103, 8.112) = 8.823, p = 0.000, Greenhouse-Geisser [GG] corrected for non-sphericity with ε = 0.276). Planned within-subjects contrasts showed significant effects for the linear (F(1, 8) = 40.869, p< 0.000) and quadratic contrasts (F(1, 8) = 7.808, p< 0.023). There was no effect of the imagery class (F(1, 8) = 1.728, p = 0.225), nor an interaction between imagery class and number of training runs (F(1.142, 9.133) = 0.423, p = 0.558, GG corrected for non-sphericity with ε = 0.285).

**Figure 2:**
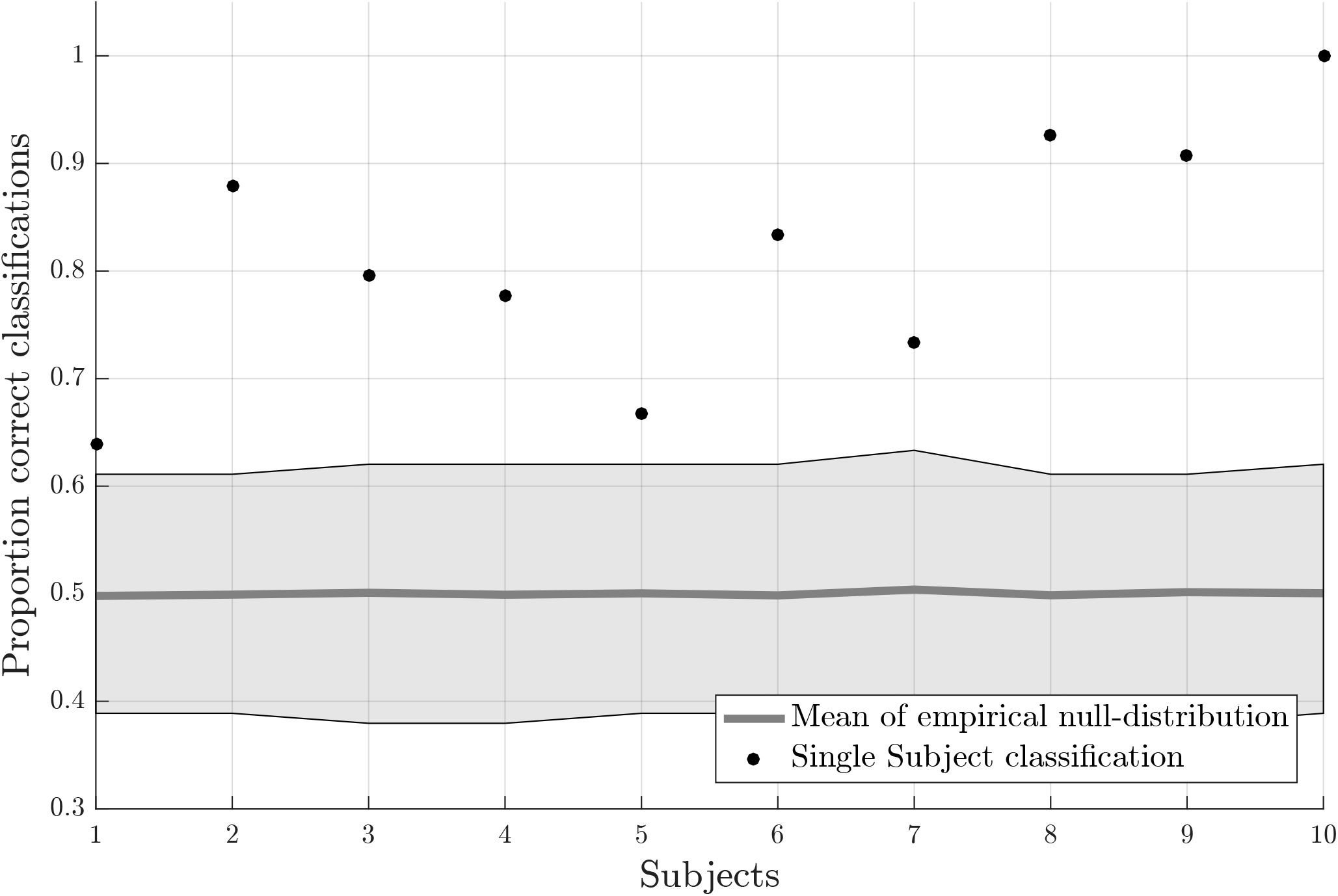
Individual offline classification accuracies. Individual single-trial classification accuracies were obtained using a leave-one-run out cross-validation scheme based on spatial features extracted from the mask combining S1 and S2 at the most lenient probability threshold. The black dots indicate the average individual classification accuracies. The grey dots and shaded area respectively represent the average of the empirical null distribution and 95% confidence interval obtained by permutation testing (2000 permutations).

**Figure 3:**
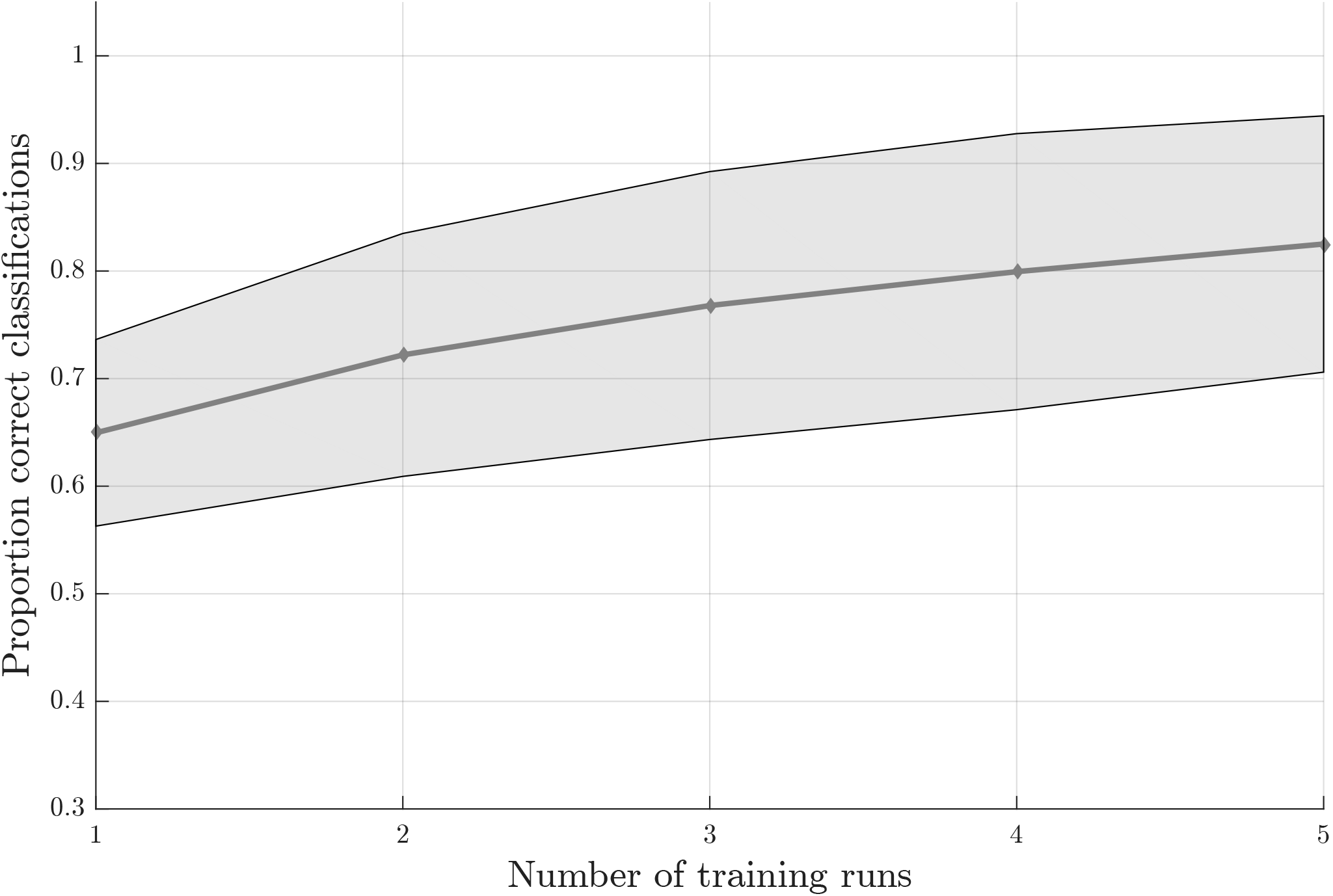
Effect of the amount of training data on the offline-obtained single-trial classification accuracy using a leave-one-run out cross-validation scheme based on spatial features extracted from the mask combining S1 and S2 at the most lenient probability threshold. The dots indicate the average classification accuracies across the nine participants who completed a total of six training runs and the shaded area indicates the standard deviation across participants.

To test the second hypothesis (1b) and provide a proof-of-concept for brain-based communication, we used the answer runs that were available from seven subjects (17 runs in total, 7 ‘no’, 10 ‘yes’; table 1). A simulated real-time analysis revealed no significant effect of the number of runs used for training (F(5, 30) = 1.470, p = 0.229, figure 4). Using all training runs, the average proportion correct for these seven subjects was 0.86 (sd = 0.17), mounting to 0.92 (sd =0.08) when disregarding one subject who, after the session, indicated he misunderstood the instructions for the answer runs. In the proof-of-concept online communication session with the last subject all answers to the questions were accurately decoded.

**Table 1:**
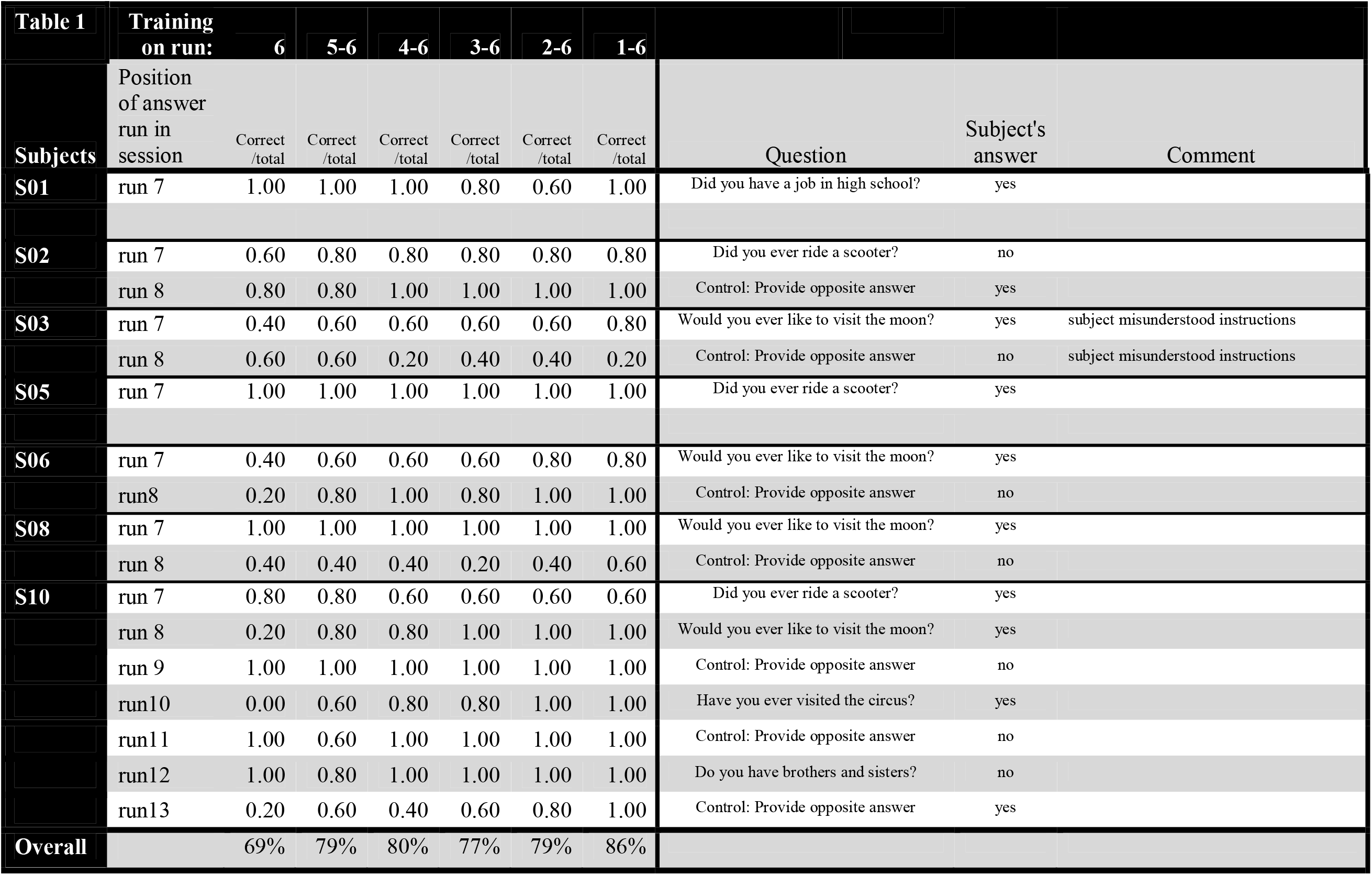
Data from simulated real-time analysis of answer runs. No answer-encoding data were obtained for subjects 4, 7 and 9 due to running short of MR scanning time.

**Figure 4:**
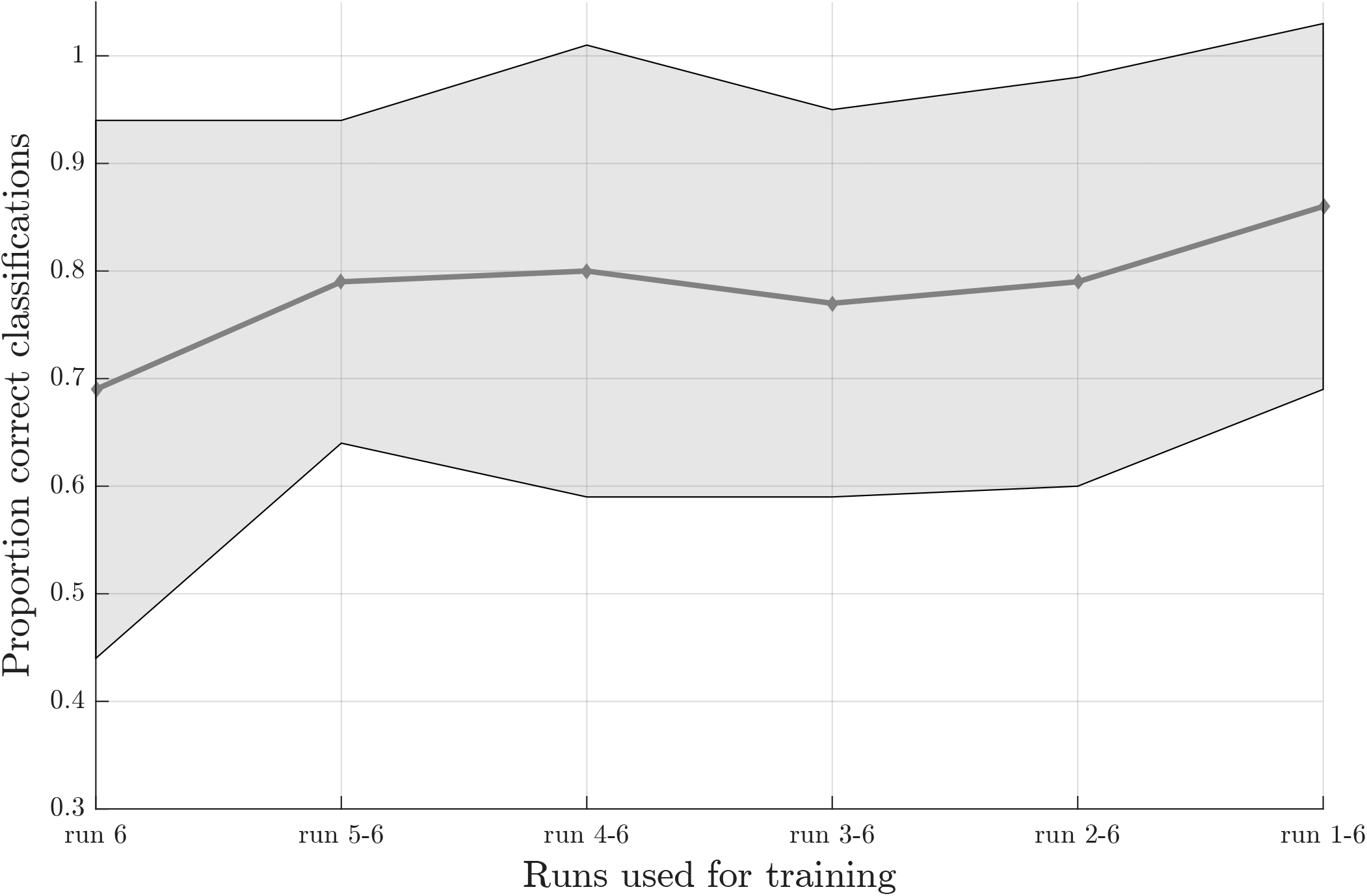
Effect of the number of training runs on average classification success in the simulated realtime procedure. The dots indicate the average classification accuracy across all participants who completed one or more answer runs and the shaded area the standard deviation.

### Assessment of somatotopy

Secondly, we wanted to assess whether classification success would be based on somatotopic information. We hypothesized (2a) that classification accuracy would be higher in primary somatosensory cortex, where somatotopy is most pronounced, compared to secondary somatosensory cortex and (2b) that the discriminative weights obtained from the support vector machine (SVM) would show a somatotopic distribution.

To test the first hypothesis (2a), we compared results obtained from anatomical probability masks of S1 versus S2 (Eickhoff, Amunts, Mohlberg, & Zilles, 2006; Eickhoff, Schleicher, Zilles, & Amunts, 2006; Geyer et al., 2001; Geyer, Schleicher, & Zilles, 1999; Geyer, Schormann, Mohlberg, & Zilles, 2000). The average proportion of correct classifications for a leave-one-run-out classification (training on all runs minus one and testing on the remaining run) was significantly higher in primary somatosensory cortex compared to the secondary somatosensory cortex (F(1, 8) = 10.581, p = 0.012, figure 5). The probability level also significantly affected classification accuracy (F(2.382, 19.060) = 20.174, p = 0.000, GG corrected with ε = .340). Decoding accuracy increased with decreasing probability level (i.e. larger mask sizes, supplementary table 2 and 3). The linear (F(1, 8) = 48.685, p = 0.000) and quadratic (F(1, 8) = 10.024, p = 0.013) within-subjects contrasts were significant. The region by level interaction was not significant (F(2.734, 21.873) = .593, p = 0.611, GG corrected with ε = 0.391). Significant above-chance classification accuracy (p<0.05) was achieved in all ten subjects at the lowest probability level (largest mask) in S1, versus seven out of ten subjects at the lowest probability level in S2.

**Figure 5:**
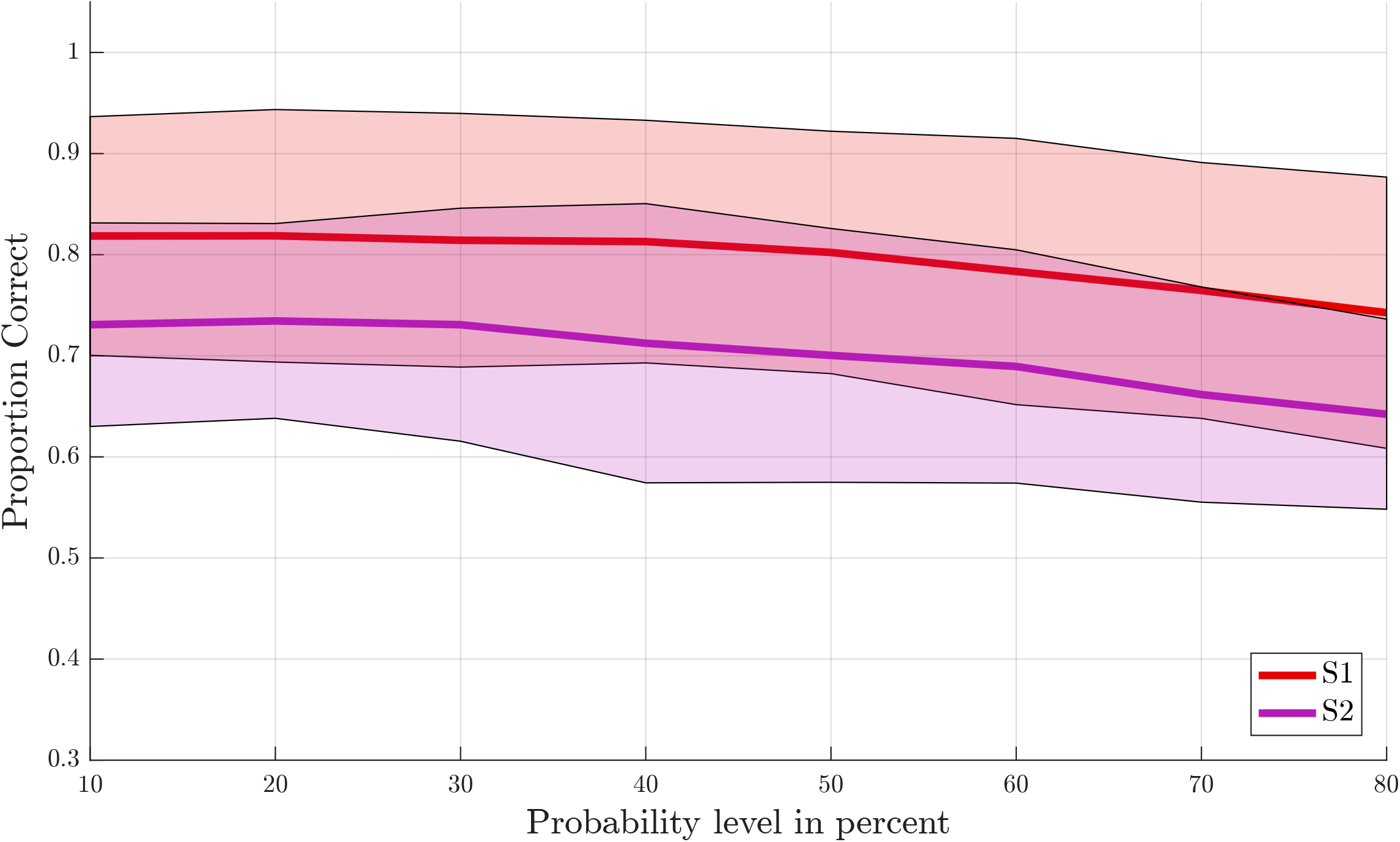
Offline single-trial classification accuracies for S1 versus S2. Accuracies were obtained using a leave-one-run out cross-validation scheme based on spatial features extracted from separate masks of S1 (red line) and S2 (purple line) at different probability levels ranging from 10% to 80%. Higher probability levels were excluded because the masks included less than 500 voxels for the S2 mask. The shaded areas indicate the standard deviation for each case.

Next, we tested somatotopy of the discriminative SVM weights (hypothesis 2b). Visual inspection of discriminative SVM weight maps revealed a pattern that roughly corresponded to the expected somatotopy in four out of ten subjects (2, 8, 9 and 10; figure 6). A weaker somatotopic pattern was observed in one subject (4). A reversed pattern was noticed in two subjects (3 and 6), and a mixed pattern was seen in three final subjects (1, 5 and 7). In a more quantitative evaluation of the extent to which there was a somatotopic distribution in the discriminative SVM weights, we tested whether there was a distinction in the median coordinates of the voxels associated with the 200 largest positive (foot) weights and the largest negative (hand) weights. We focused on the x-coordinates only, because somatotopy is expected to be most pronounced on the medial-lateral axis of the brain. A Wilcoxon signed rank test revealed that there was a significant within-subject difference (z = 2.09, p = 0.0365) between the median x-coordinates of the voxels associated with the 200 largest positive (foot) weights and the median x-coordinate of the voxels associated with the 200 largest negative (hand) weights. The median x-coordinate across subjects was 8.50 (sd = 14.5) for the 200 largest positive weights and −21.5 (sd = 18.4) for the 200 largest negative weights (figure 7). Also, we found a significant correlation between the difference in median × coordinates and the overall classification accuracy, i.e. subjects with a larger value for the foot-hand median x-coordinate difference (larger ‘hand-foot distance’) also showed a higher classification accuracy (Spearmann’s rho .948, p = 0.000).

**Figure 6:**
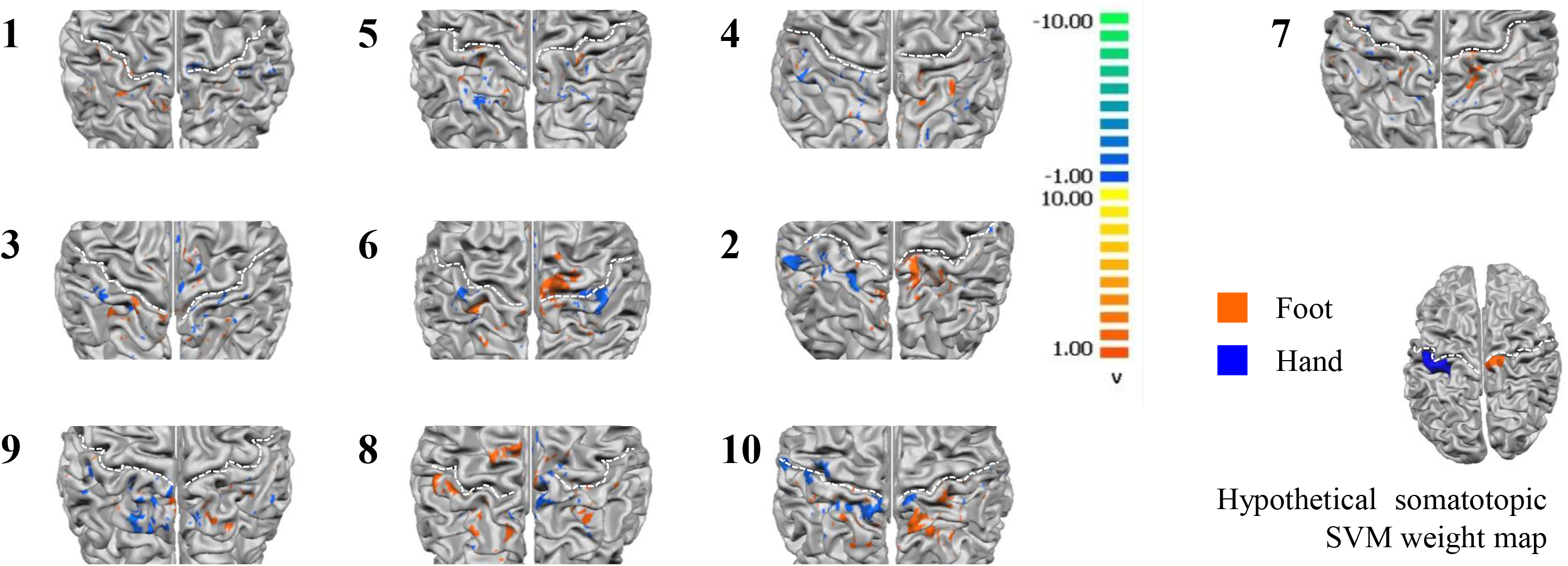
Z-scored average individual weight maps obtained from the combined S1 and S2 mask at the most lenient probability threshold, projected on individual cortical surface reconstructions. Individual subject data were ordered by average classification accuracy in the leave-one-run out cross validation procedure (accuracy increases from left to right and top to bottom). The map of participant 7 is shown separately, because for this, only four training runs could be used for testing instead of the five available for the other participants. Warm colors indicate positive weights (one could say voting for “foot”), cold colors for negative weights (a vote for “hand”). The white dotted lines indicate the central sulci. Hypothetical somatotopic SVM weight pattern with similar color coding is shown at the bottom right of the figure, based on the approximate location reported for the hand and foot representations in the primary somatosensory cortex. Note that the foot representation extends onto the medial surface.

**Figure 7:**
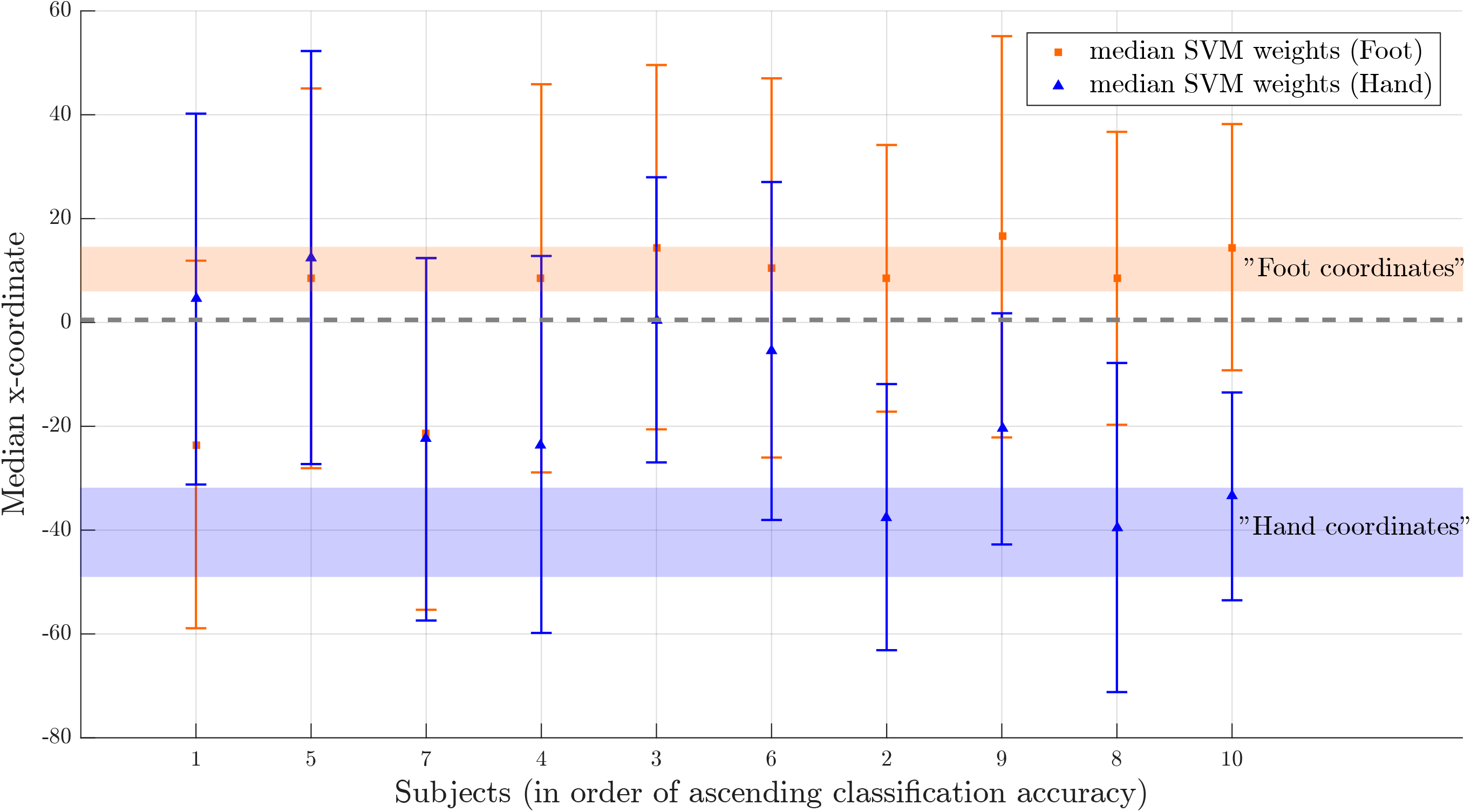
Weight distribution on the medial-lateral brain axis for hand and foot imagery. Median x-coordinate of the 200 largest positive and negative weights for individual subjects, ordered from left to right according to increasing classification accuracy. The shaded region indicates expected location for foot (orange) and hand (blue) on somatotopic map based on values reported in the literature (Akselrod et al., 2017; Nelson & Chen, 2008). The dotted line indicates the brain midline.

## Discussion

The current study (1) investigated whether somatosensory imagery is a suitable encoding strategy for a hemodynamic (fMRI-based) BCI, aiming to provide a proof-of-concept of its suitability for online BCI-based communication and (2) tested to what extent its decoding is based on somatotopic information. In accordance with our first set of hypotheses, we found that the contents of somatosensory imagery could be decoded above chance in each of our participants with similar accuracy for hand or foot, and that decoding accuracy increased with an increasing number of training runs. We also found that answers to binary questions encoded by seven healthy participants using the somatosensory imagery BCI could be successfully decoded in simulated real-time as well as during a proof-of-concept online communication session tested in one subject. In accordance with our second set of hypotheses, we found that decoding accuracy was significantly higher in primary somatosensory cortex compared to secondary somatosensory cortex, and that discriminative weight patterns generated by healthy participants through somatosensory mental imagery showed a somatotopic distribution.

### Suitability for use in fMRI Brain-Computer-interfacing

The average classification success was 82% for the offline leave-one-run out cross-validation procedure on the training runs, and 92% for the classification of the answer runs in simulated realtime (86% when including a subject who indicated afterwards that he misunderstood the instructions). The fact that the classification success was higher for simulated real-time classification of the answer runs could be explained by the additional training run that was used for training the classifier in simulated real-time, whereas in the offline leave-one-run out procedure one run necessarily had to be left out for testing. However, the evaluation of the effect of additional training runs in the offline condition suggested that the beneficial effect of adding training runs levelled off after four runs. Another possible explanation might lie in the slight differences in the analysis pipelines for the simulated real-time and offline analyses regarding slice-scan-time-correction and data transformation (Talairach transformation and downsampling to 2mm^3^ offline versus smoothing with 4mm FWHM Gaussian kernel in native space in simulated-real-time). On the other hand, the increased accuracy for the answer run classification might also be related to enhanced imagery quality during the answer run, due to extended practice, focus on a single imagery strategy (in contrast to the alternating imagery strategies in the training runs) and possibly increased motivation because of the social interaction component of answering a genuine question.

Taken together, these findings inspire confidence that somatosensory imagery will be a viable additional control strategy for BCI communication procedures. Somatosensory imagery might be especially suited for locked-in patients with reduced visual or auditory faculties who lack muscle control. These patients might not be able to evoke stable imagery involving their most affected modalities, as they lack recent memories of sensory experiences which can increase the level of detail and vividness of mental representations generated by imagination (Hassabis & Maguire, 2009). The current study was performed at 7T, but given the relatively large distance between hand and foot representations in primary somatosensory cortex similar results could be expected at a field strength of 3T which is more commonly available in clinical settings. Our novel somatosensory imagery control strategy might also be applied in binary fNIRS-based communication BCIs. Of course, taking into account the considerably lower spatial resolution of fNIRS, optode placement and data analysis procedures would have to be optimized.

### Topographic nature of somatosensory imagery

Our results indicate that decoding accuracy was higher within the primary than in the secondary somatosensory cortex mask, and that it is possible to induce somatotopic discriminative weight patterns by mental imagery. The median x-coordinates of the subjects with highest offline classification accuracies roughly correspond to values reported in the literature for finger (Kolasinski et al., 2016; Nelson & Chen, 2008; Pfannmoller, Greiner, Balasubramanian, & Lotze, 2016; Sanchez-Panchuelo, Francis, Bowtell, & Schluppeck, 2010; Schweizer, Voit, & Frahm, 2008) and foot representations (Akselrod et al., 2017; figure 7). The slightly more medial imagery-related weight coordinates could be due to the fact that, in the current study, imagery involved the whole hand whereas stimulation is usually delivered to the fingers only. For the foot, imagery coordinates were more inferior than reported stimulation coordinates. Similar observations of corresponding sites for imagery and stimulation in primary sensory cortices have also been made in previous studies in the somatosensory domain (de Borst & de Gelder, 2017; Newman et al., 2005; Schmidt et al., 2014; Wise et al., 2016; Yoo et al., 2003), however, to our knowledge, the current study is the first to report a somatotopic pattern of discriminative weights in the (primary) somatosensory cortex during imagery. Previous studies have reported topographic recruitment of visual cortex during imagery (Klein et al., 2004; Slotnick, Thompson, & Kosslyn, 2005; Thirion et al., 2006)

The presence of a somatotopic pattern in the discriminative weights, that is a more medial and right sided location for the highest positive weights corresponding to the imagery class ‘foot’ and a more lateral and left sided location for the highest negative weights associated with ‘hand’, was associated with higher classification success. This suggests that subjects might increase classification success by increasing the level of somatotopy in their imagery-induced brain activation patterns. If this finding can be extrapolated to subjects with maladaptive somatosensory representations, we speculate that the current BCI might be suitable to use for training the somatotopy of these representations. Somatosensory dysfunction and aberrant plasticity have been linked to pain severity in chronic pain (Di Pietro et al., 2013; Di Pietro, Stanton, Moseley, Lotze, & McAuley, 2015; Flor, Braun, Elbert, & Birbaumer, 1997; Kim, Kim, & Nabekura, 2017; Pleger et al., 2005; Wrigley et al., 2009), but see Makin et al. (2013) for an alternative view). Neurofeedback based on somatosensory mental imagery could be especially suitable for clinical interventions in patients for whom actual somatosensory stimulation is difficult or impossible due to their physical limitations or technical limitations of the scanner environment.

Somatotopic patterns could in theory also be evoked using motor imagery or even actual movement. However, our subjects were carefully instructed and trained in a mock scanner to exclusively use somatosensory imagery, and it was emphasized that they should not move during imagery runs. Subjects’ compliance to these instructions is supported by the fact that we did not observe hand or foot movement during training or scanning. Also, during the training runs, subjects did not receive any feedback or reward based on their performance, so there was no incentive to try out non-somatosensory imagery strategies. In line with this, subjects’ reports indeed indicated that they used purely somatosensory strategies (supplementary table 1).

Based on the current data, we cannot establish a link between classification accuracy and the use of a specific somatosensory imagery strategy: The two subjects with largest classification success used touch and warmth imagery, as did the two subjects with the lowest classification success. It would be interesting to investigate whether imagery targeting different somatosensory submodalities induces different patterns of activation in primary somatosensory cortex. However, such pattern differences might be most apparent within a limb-specific subregion, whereas the general somatotopy of hand and foot is most likely preserved across somatosensory submodalities (Sur, Wall, & Kaas, 1984).

We instructed our subjects to stick with the same strategy for hand and foot. Whereas some subjects only reported using imagery based on one somatosensory submodality (e.g. vibrotactile stimulation), others used imagery of more naturalistic somatosensory stimulation involving a combination of somatosensory submodalities. Such imagery, e.g., combining warmth and touch might increase vividness and evoke stronger activation. Work by Kosslyn and Thompson in the visual domain suggests that the level of vividness and detail is related to primary visual cortex activation during imagery (Kosslyn & Thompson, 2003). We therefore propose to leave the choice of strategy to the participant, and have them choose the strategy that subjectively evokes most vivid and detailed imagery and that they feel most comfortable with.

There are some limitations to this study. Somatosensory imagery involved only two contralateral body parts, which are relatively far apart in physical and somatotopic space, and whose representations reside on contralateral sides of the body. Thus, classification could have been based solely on less specific imagery related to body side. However, the fact that classification success was highest within the primary somatosensory cortex mask and linked to the degree of somatotopy in the discriminative weight maps indicates that subjects used a limb-specific somatosensory imagery strategy rather than a body-side related strategy. This is also in accordance with the introspective description of imagery strategies reported by the subjects. However, note that the question whether the brain signal is body-part or body-side specific is irrelevant in the context of the suggested two-choice communication BCI application, as long as it can be used at will.

A final limitation of this study is that only healthy subjects without neurological impairments participated in the current study. Patients might have shorter attentional spans leading to more variable imagery quality. However, we found higher classification rates during answer runs than during training runs, possibly due to increased motivation. This effect might positively extrapolate to patients in a BCI communication setting.

### Conclusion

Taken together, we conclude that somatosensory imagery is a successful information-encoding strategy for hemodynamic BCI-based communication with a high clinical potential. The current results pave the way for future applications, such as brain-based communication for patients with locked-in-syndrome, and neurofeedback-based training to support the rehabilitation of somatosensory representations in stroke and chronic pain patients suffering from somatosensory deficits.

## Acknowledgements

This work was supported by the *Braingain SmartMix* grant (A.L.K., B.S. and G.V.) and *European Research Council* (ERC) Grant 269853 (R.G.). This work was also supported by the *European Commission* (7th Framework Programme 2007-2013, DECODER project, B.S and R.G.). We thank all participants and the students, Mona Rosenke, Tobiasz Kaduk and Cynthia van de Wauw, who helped to acquire and analyze the data presented in this paper.

