## Supplementary Tables for "Topographic somatosensory imagery for real-time fMRI Brain-Computer Interfacing"

Supplementary table 1: Data from exit questionnaire on imagery strategies

**Supplementary Table 1**

*Data from exit questionnaire on imagery strategies*

| | Quality of imagery (0-10) | | Tactile sensation (0-10) | | % of time imagery worked | | Description of imagery | |
|---|---|---|---|---|---|---|---|---|
| *Subject* | *hand* | *foot* | *hand* | *foot* | *hand* | *foot* | *hand* | *foot* |
| S01 | 7 | 6 | 3 | 2 | 79 | 64 | Warm touch remembered from real touch | Same as hand but translated to foot |
| S05 | 7 | 7 | 8 | 8 | * | * | I tried to imagine touch | I tried to imagine touch |
| S07 | 7 | 6 | 2 | 3 | * | * | I mostly directed attention to the respective spot and tried to imagine the sensation | I mostly directed attention to the respective spot and tried to imagine the sensation |
| S04 | 6 | 6 | 4 | 5 | * | * | I tried to imagine stimulation from the piezo stimulator during the imagination | I tried to imagine stimulation from the piezo stimulator during the imagination |
| S03 | 8 | 7 | 8 | 6 | 70 | 80 | Hot feeling across the hand | Hot feeling across the foot |
| S06 | 7 | 7 | 5 | 5 | * | * | My strategy was to imagine pinpricks similar to the piezo stimulation on my hand and foot (although this was more difficult on the foot) | My strategy was to imagine pinpricks similar to the piezo stimulation on my hand and foot (although this was more difficult on the foot) |
| S02 | 7 | 7 | 7 | 7 | 75 | 75 | Imagine like warm water flowing all around my hand | Imagine like warm water flowing all around foot |
| S09 | 6 | 7 | 4 | 3 | 69 | 58 | Paint-brush on top of hand stroking each finger one at a time, starting from lateral side moving medially | Paint-brush stroke on soleof foot-starting from lateral side moving up and down towards medial side |
| S08 | 8 | 6 | 8 | 6 | 94 | 84 | heat (mainly in the palm of my hand) | heat (mainly in the middle part of the sole of foot) |
| S10 | 6 | 7 | 4 | 4 | 98 | 98 | I imagined being massaged with strong circular thumb finger movement on the inner side of my palm. | I imagined being massaged with strong circular thumb finger movement on the inner (soft) side of my foot. |

** data missing due to a mistake*

Supplementary table 2: Number of functional voxels for each anatomical region and probability level.

| **Supplementary Table 2** | | |
|---|---|---|
| *Number of functional voxels for each anatomical region and probability level* | | |
| | Area | |
| Probability Level (%) | S1 | S2 |
| 100 | 451 | 5 |
| 90 | 1031 | 169 |
| 80 | 1701 | 500 |
| 70 | 2556 | 917 |
| 60 | 3721 | 1444 |
| 50 | 4986 | 2124 |
| 40 | 6373 | 2816 |
| 30 | 8232 | 3673 |
| 20 | 10638 | 5011 |
| 10 | 14543 | 7254 |

Supplementary table 3: Average decoding accuracy for each anatomical region and probability level based on cross validation with N-1 training runs and 1 test run.

**Supplementary Table 3**

*Average decoding accuracy for each anatomical region and probability level based on crossvalidation with N-1 training runs and 1 test run.*

| | Areas | |
|---|---|---|
| Probability Level (%) | S1 | S2 |
| 100 | 0.67 | 0.51 |
| 90 | 0.73 | 0.60 |
| 80 | 0.74 | 0.64 |
| 70 | 0.76 | 0.66 |
| 60 | 0.78 | 0.69 |
| 50 | 0.80 | 0.70 |
| 40 | 0.81 | 0.71 |
| 30 | 0.81 | 0.73 |
| 20 | 0.82 | 0.73 |
| 10 | 0.82 | 0.73 |
